# Retinal waves reveal axial biases in modular patterns of cortical activity that predict future orientation preferences

**DOI:** 10.1101/2025.07.09.663735

**Authors:** Alexandra Gribizis, David Fitzpatrick

## Abstract

Spontaneous activity before sensory onset is thought to guide the formation of functional neural circuits. In visual cortex, spontaneous activity prior to experience exhibits modular patterns that resemble future visually evoked orientation selective responses. However, the factors at this early stage that build the initial network interactions that support the orientation tuning of modular responses at eye opening remain unknown. Here we provide the first evidence that retinal waves could play an important role by shaping the modular biases in patterns of intracortical connectivity and lay the foundation for orientation selective responses. We demonstrate that slow propagating waves in developing cortex that are dependent on retinal activity recruit specific modular patterns during their movement across the cortical surface, resulting in strikingly elongated patterns of modular coactivity that predict visually-evoked orientation responses at eye opening. Thus axial biases in spontaneous patterns of co-activity are present before the onset of visual experience and could serve as the seed for the developmental emergence of modular orientation representations.

## Introduction

Developing neural circuits exhibit spontaneous patterns of activity that prepare the brain before sensory onset^1–5^. The mature visual cortex in highly visual animals is characterized by iterated clusters of neurons, or modules, that preferentially respond to similar stimulus features, such as orientation. Previous work has shown that before the onset of visual experience, spontaneous activity in the cortex can exhibit modular patterns that resemble future orientation map structures^6^, an initial framework that visual experience then strengthens and builds upon^7^. At the same time, early development of the visual system is also characterized by retinal waves—linear wavefronts of spontaneous activity that propagate across the retina and through downstream visual pathways prior to eye opening^8,9^. These retinal waves are thought to play a critical role in refining retinotopic maps but are considered largely independent from the modular, intracortical activity patterns that emerge around the same developmental window. As a result, the potential interaction between spontaneous retinal activity and the developing intracortical modular network has remained largely unexplored.

In the mature primary visual cortex (V1), long-range horizontal axonal connections span several millimeters and preferentially link columns with similar orientation preferences. Rather than spreading isotropically, these connections tend to follow the axis of preferred orientation, forming structured networks that are thought to support perceptual processes such as contour integration by linking co-oriented and coaxially aligned receptive fields^10^. In the present study, we investigate how early spontaneous activity driven by retinal waves may contribute to the organization of these intracortical networks. Using chronic calcium imaging in developing tree shrews—a species with a well-defined modular visual cortex— we track spontaneous cortical activity both before and after eye opening. Our findings reveal, for the first time, the presence of retinal-driven cortical waves with a modular structure that is specific to the topographic axis of the wavefront. Strikingly, we observe a high degree of correspondence between the axial biases in modular coactivity driven by these early spontaneous cortical wavefronts and the orientation preference maps that emerge after visual experience begins. These results suggest that the structural and functional organization of orientation networks in the cortex may be strongly shaped by retinal activity well before the onset of vision.

## Results

### Spontaneous retinal-driven waves recruit emerging modular networks in developing visual cortex

To investigate the relationship between spontaneous retinal wave activity and the developing cortical modular network, we chronically recorded cortical activity in neonatal tree shrews by first injecting viral GCaMP7s into V1 of early postnatal shrews (postnatal days 2-4, p2-p4, Fig 1A). A custom designed metal head plate 8mm in diameter was then implanted over the injected region after 10-12 days of expression, and then imaging was conducted on subsequent days up until eye opening. All shrews were allowed to naturally open their eyes ranging from postnatal day 18 to postnatal day 21. In a subset of shrews (3), we also imaged several days past eye opening (p23+) to directly compare visually-evoked responses to oriented stimuli with the spontaneous activity before eye opening.

**Figure 1:**
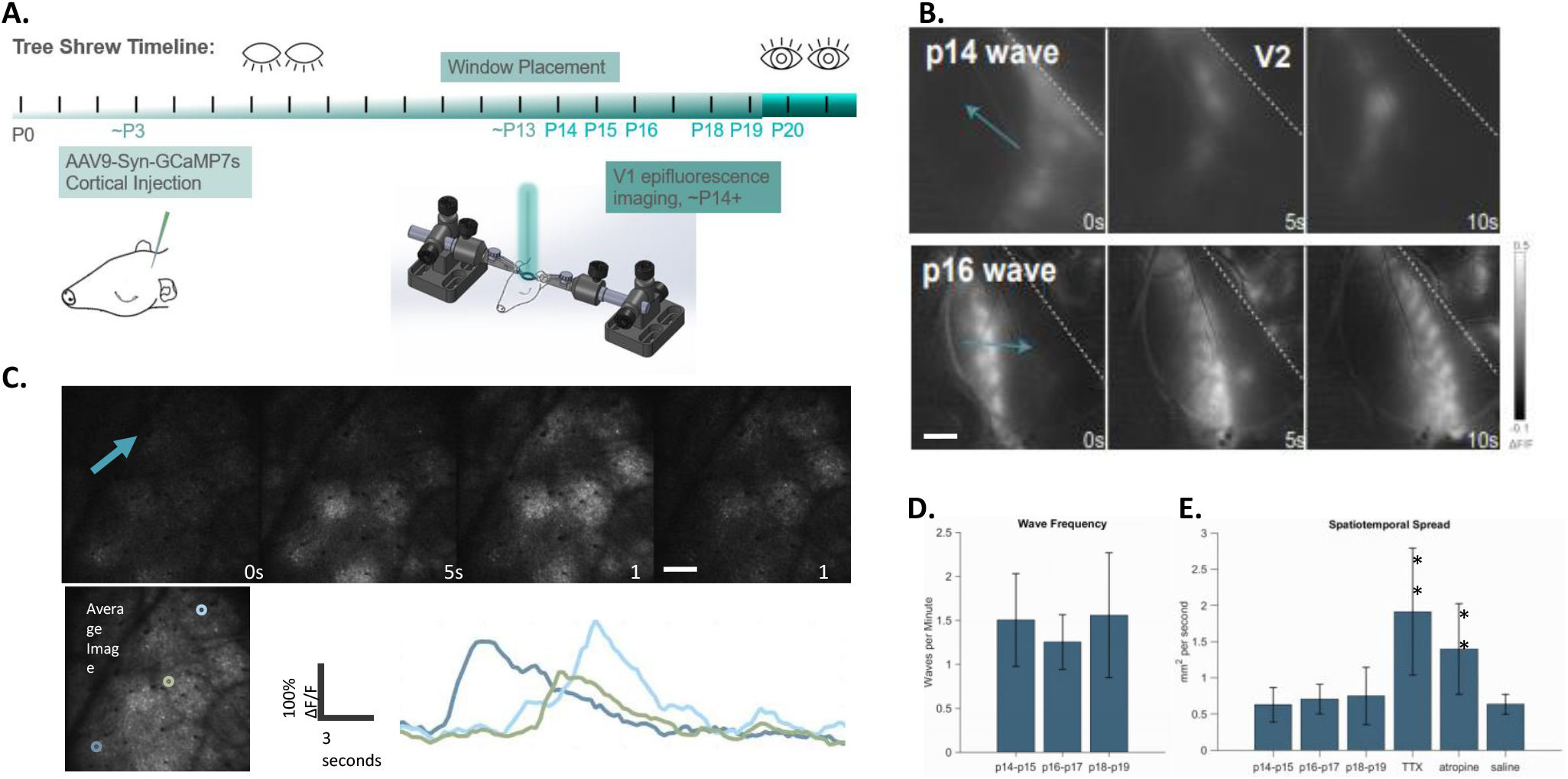
Spontaneous retinal-driven waves recruit emerging modular networks in developing visual cortex. **A**. Graphical depiction of experimental timeline. P=postnatal day. **B**. Chronic widefield epifluorescence calcium imaging of spontaneous activity at 2 early age timepoints capturing spontaneous waves traveling across V1, and the emergence of modular patterns in the P 16 waves. Scale bar=1mm. **C. (top)** 2-photon imaging of cell groups during a typical spontaneous wave on postnatal day 16. Scale bar 500 um. **(bottom)** Sequential activity of 3 individual cortical neurons at different cortical locations during the wave. Color coded location of cells on the left, individual cell traces on the right. **D**. Frequency of wave activity the week before eye opening measured at the pixel level, i.e. waves across each pixel location during recording. Wave frequency remains between 1-2 waves per minute across ages in the week before eye opening. (n=5 tree shrews. Error bars display standard deviation). **E**. Spatiotemporal spread of spontaneous propagating waves across age groups shown in Fig. 1D and conditions where either TTX or saline were injected into the eye during postnatal day 17 recordings (n=3), or atropine (dose) was provided subcutaneously (atropine experiments, n=3, TTX and saline experiments, n=2). ** = p<0.01. One way ANOVA with correction for multiple comparisons.

In the earliest recordings (Fig 1B top), ∼p14, we saw evidence of large, slow propagating waves across the V1 surface; later recordings in the same shrew (Fig 1B bottom) demonstrated that slow propagating waves had acquired a modular recruitment pattern that was also evident in cellular scale 2-photon imaging (Fig 1C). Wave frequency across each pixel location was constant through each age group (Fig 1D: p14-p15, average 1.5 waves/minute, standard deviation of 0.5; p16-p17, average 1.2 waves/minute, standard deviation 0.3; p18-p19 average 1.5 waves/minute, standard deviation 0.7, n=5 chronically imaged tree shrews, no significant difference between groups measured by one-way ANOVA), consistent with the frequency of retinal waves^8,9,11–13^. The slow spatiotemporal spread that defines the wave activity (see methods) was also constant between age groups (0.62 +/- 0.24 mm^2^/second, 0.7 +/- 0.2 mm^2^/second, 0.74 +/- 0.4 mm^2^/second respectively for p14-p15, p16-p27, and p18-p19 age groups, Fig 1E). In 3 separate acute experiments at postnatal day 17, tetrodotoxin (TTX) was injected into the vitreous of the contralateral eye to remove retinal wave activity^8,14^. After confirming the removal of retinal activity with the absence of calcium response to flashing stimuli, subsequent recording revealed large activations remained in visual cortex as had been previously described in ferret^6^, but slow propagating waves had been eliminated (spatiotemporal spread 1.9 +/- 0.87 mm^2^/second, saline control before TTX injection 0.63 +/-0.14). This elimination of slow propagating waves was also evident 30 minutes after subcutaneous atropine injection in the chronically imaged tree shrews (1.4 +/- 0.63 mm^2^/second, p<0.01. One-way ANOVA with correction for multiple comparisons) which has been shown to block wave propagation^15,16^.

### Specific modular patterns recruited based on the topographic axis of the wavefront

To understand the relationship between waves and the modular patterns they evoke, we first examined 2 waves propagating in approximately orthogonal directions (Fig 2A top), and compared the total pattern over the duration of each wave (spatial Pearson correlation, r=0.12). The distinct modular patterns associated with these two different wave directions was evident both in the color-coded depiction of the sum of the two waves and in the modular pattern resulting from the difference in activity driven by the two wave directions (Fig 2A bottom). To test the consistency in the relationship between the topography of wave motion and the identity of the modules that are activated we determined the broad patterns of modular activity across the cortical surface associated with different seed points. Indeed, seed-correlations (see Methods) of an individual module derived from an entire recording session revealed robust modular network patterns that were specific to the seed point. To test the specificity of the patterns, we compared modular correlation patterns found for nearby seed points that were located in domains with non-correlated (NC) vs correlated patterns of spontaneous activity. The patterns of modular activity associated with NC seed points were not only distinct in their modular distribution (Fig 2B far right), but also in their axis of elongation across the cortical surface as can be appreciated by comparing the correlation pattern distributions with the V1-V2 border (Fig 2B left and middle). In contrast, and as expected, modular correlation patterns for nearby seeds within domains with correlated spontaneous activity (C) had a high degree of modular overlap and they also exhibited similar elongation axes (Fig 2E). The degree of modular and axial specificity of the seed point correlation patterns is quantified in Fig. 2F for n=4 tree shrews with 12 initial seeds each. For modular specificity, we calculated spatial 2D Pearson correlation between networks and found for NC network seed correlations a value of 0.02 +/- 0.02 vs. 0.49 +/- 0.12 for C network seed correlations. For axial elongation bias, the angular difference between NC seeds was 78 +/- 29 degrees while for C seeds it was 6.1 +/- 2.4 degrees). Consistent with a wave driven origin of the elongation axis, we found that the axial elongation is reduced in the presence of wave-abolishing atropine such that the width of the bounding ellipse is significantly widened (Figs 2C and 2D, *** p<<0.001, paired t-test), (see Discussion).

**Figure 2:**
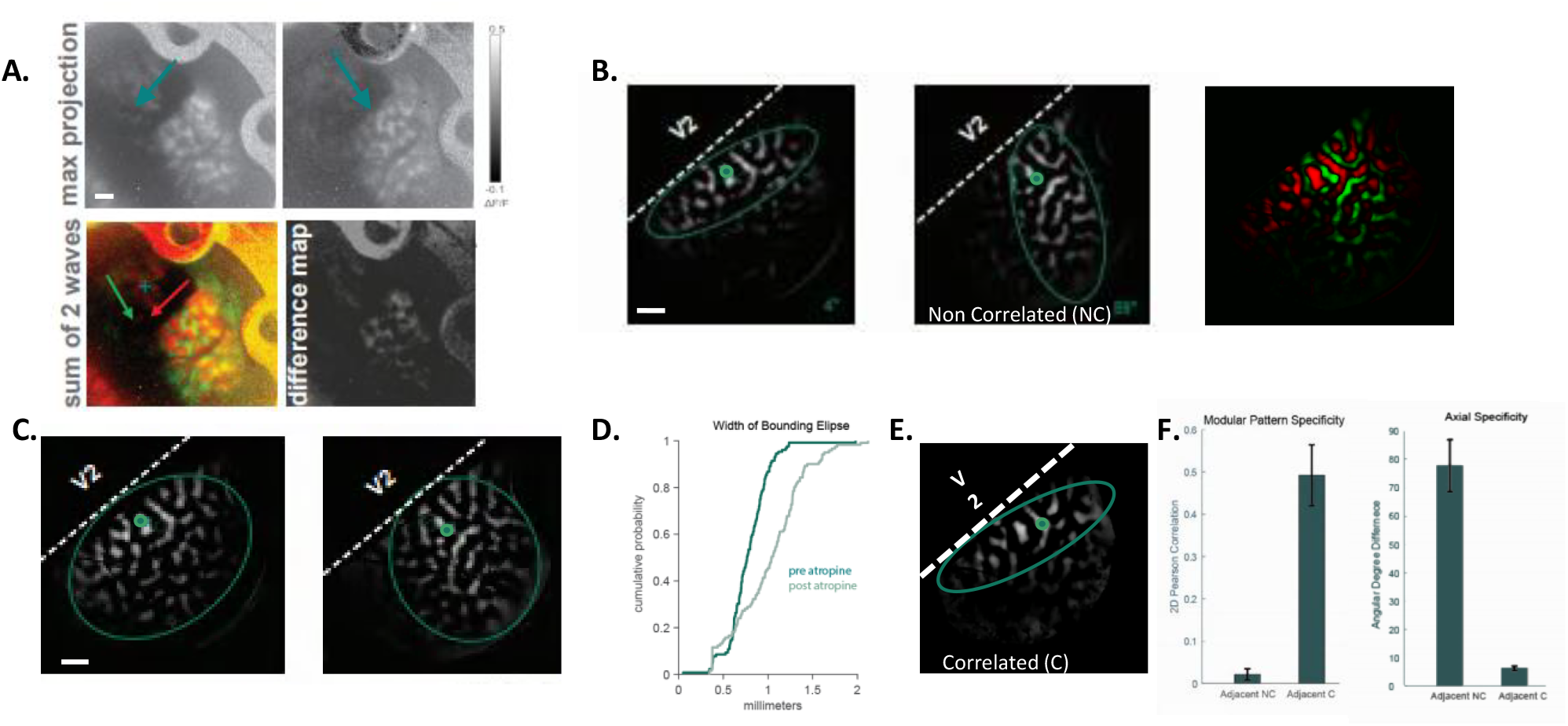
Specificity in the modular and axial patterns of activity recruited by spontaneous wavefronts. **A**. (**top**) Cumulative activity resulting from two waves moving in nearly orthogonal directions (green arrows) during P16 recording. (bottom) Complementary spatial relationships of the modular patterns associated with these waves including sum (left) and difference images (right). Scale bar = 1mm. **B**. (left) The average pattern of pixel activity correlated with one seedpoint (green dot) during P19 recording, illustrating an axial bias elongated parallel to the V1/V2 border. Scale bar = 1mm. (middle) Average pixel activity correlated with a second seedpoint (green dot) placed in a nearby region not correlated with the first seedpoint and the resulting axial bias elongated away from the V1/V2 border. (right) Overlay of the complementary modular patterns for the two seed points. **C**. Impact of blocking wave activity with subcutaneous atropine injection on the average pixel activity patterns for the 2 seed points in B. **D**. Width of ellipsoid envelope (see Methods) before and after atropine, n=4. E. Pixel correlation pattern from a third seedpoint that is within the correlation pattern of the first seedpoint, illustrating a similar axial bias. **F**. (left) Quantitative comparison of modular patterns (2D Pearson spatial correlations) for the first seedpoint with the adjacent non-correlated seedpoint (NC), and the adjacent correlated seedpoint. (right) Similar comparison for axial biases (angular degree of difference in the enveloping ellipsoids) n=4, **p<0.01 for both groups using KW nonparametric testing; p<0.01 for spatial correlation and ***p<0.001 for angular differences using one-way ANOVA.

### Axial topology of spontaneous modular patterns predicts future orientation preference

Given the specificity of modular network correlations presented with a strong axial bias that was location dependent, we next wanted to know if the relationship of the axial bias was reflective of the future orientation selectivity of a given module. To answer this question, we took advantage of the access to the V1/V2 border in our recordings, which gave us a landmark for the future map of visual space; the V1/V2 border is known to define the vertical meridian in visual space^17,18^. We related seeds that were parallel to the V1/V2 border (Fig 3A) and orthogonal to the border/vertical meridian (Fig 3B) during spontaneous activity before eye opening (p19) to visual responses after eye opening (p23). To do this, we overlaid the resulting seed-correlation during spontaneous activity before eye opening with the orientation map in the same animal after eye opening to produce an effective tuning curve of the spontaneous modular network (Figs 3C and 3D, see Methods for more detail). Predicted tuning curves were significantly associated to their orientation responses after eye opening based on their angular relationship to the vertical meridian, with stronger relationships in the cardinal responses than oblique, p<0.001 for vertical and horizontal orientations, p=0.09 for 45 degrees, p=0.02 for 135 degrees, circular V-test for predicted orientations based on angular relationship to VM, n=3 tree shrews at age p19.

**Figure 3:**
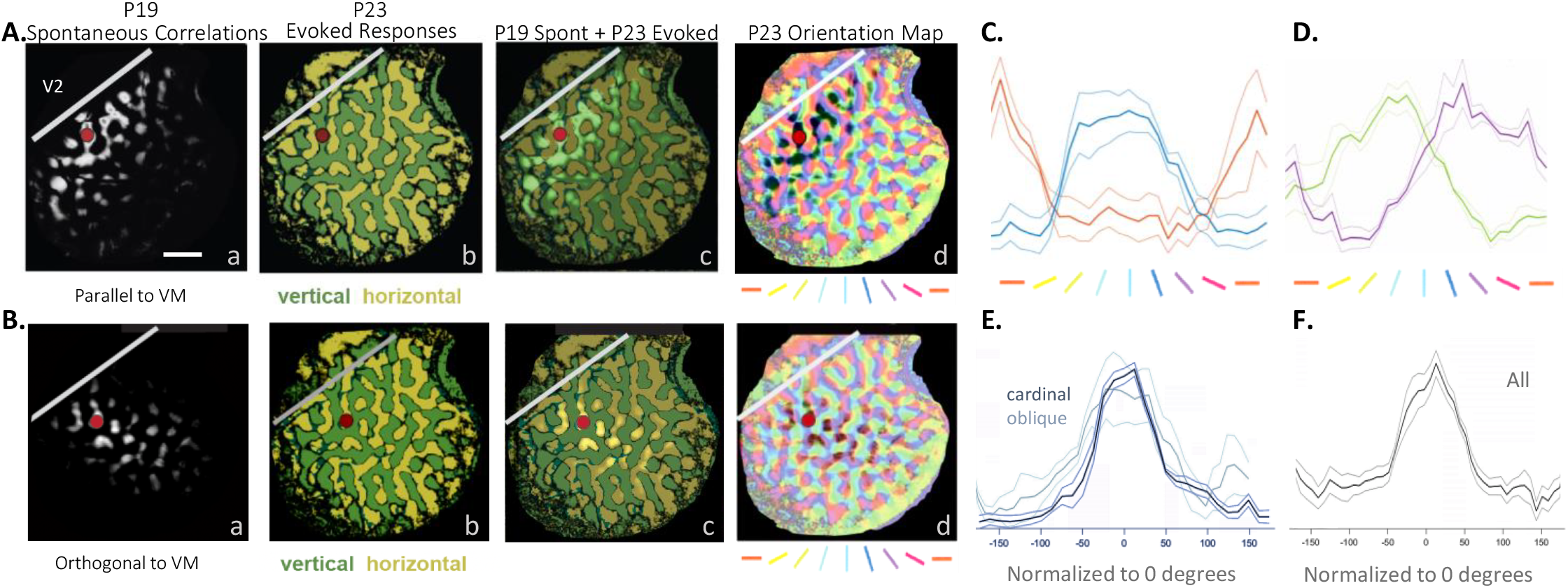
Axial topology of spontaneous modular patterns predicts future orientation preference. **A. (a)** Pixel-based correlation (see methods) of spontaneous recording at postnatal day 19 reveals axial distribution of co-active areas parallel to the vertical meridian, VM (scale bar = 1mm). **(b)** Evoked vertical (dark green) and horizontal (light green) responses in the same animal after eye opening at P23. **(c)** P19 spontaneous correlations (light green) overlaid on P23 evoked responses showing overlap predominately with vertical evoked responses (dark green). **(d)** P19 spontaneous correlations overlaid with polar map of orientation preference from P23. **B**. Same as in **A** but using P19 spontaneous correlations from a nearby location with axial alignment orthogonal to the VM. These P19 spontaneous modular patterns overlap predominantly with horizontal P23 evoked responses. **C**. Quantifying axial topology predictions of future orientation preference by overlaying P19 spontaneous patterns with axial alignment parallel and orthogonal to the V1-V2 border with the P23 evoked orientation preference map. Normalized predicted tuning curves for parallel and orthogonal elongation axes were well correlated with the evoked representations of vertical and horizontal gratings respectively p<0.001; circular V-test for predicted orientations based on angular relationship to VM, n=3 tree shrews. **D**. Similar predictions for oblique P19 spontaneous axial biases (45 degrees, p=0.09; 135 degrees p=0.02. **E**. Cardinal or oblique orientations normalized to zero degrees. **F**. All predictions of future orientation preference normalized to zero degrees.

## Discussion

Here we present data from chronically imaged neonatal tree shrews demonstrating that spontaneous retinal-driven waves in visual cortex exhibit a modular structure in a species with a columnar architecture. These waves recruit specific neuronal populations aligned with the wavefront’s topographic axis and form elongated, long-range correlated networks. Most interestingly, the axial bias of the correlated networks for a given module predicts the future orientation preference after eye opening. This relationship is consistent with the long-range orientation-specific bias in horizontal connections that align with cells preferred orientation in adult tree shrew^10^. This is the first study that shows a relationship between spontaneous retinal waves and modular structure in developing cortex, and that these early waves drive specific patterns of modular activity that are consistent with visually driven orientation responses following the onset of visual experience.

While previous developmental studies have primarily focused on rodents and carnivores, species with delayed eye opening (approximately p14 in mice and p30 in ferrets^19^), this is the first study examining developing visual cortex in the tree shrew. The mature tree shrew has also long served as a model for studying the functional organization of mature circuits in V1, due to its well-defined modular representation of orientation and resemblance to primate neocortex^20,21^. Tree shrews, like mice, have a smooth cortex that is unencumbered by shifting sulci during rapid growth and are also born approximately 20 days before eye opening. Unlike tree shrews, *murine* species have a “salt and pepper” organization of orientation preference, albeit with feature-specific microcircuits^22^. Recent recordings from other developing columnar species^23,24^ have not reported slow propagating modular waves of confirmed retinal origin, possibly due to limitations in targeting and accessing V1 for imaging studies, or recording before modular structure is likely to have formed^24^ (such as p14 in tree shrew, Fig. 1B). The visualization of modular wave activity was fundamental to the novel findings in this study since it revealed a preferred elongation axis that made it possible to relate spontaneous modular activity to future visually evoked orientation responses, an axis that was not observable after the removal of wave activity (Fig. 2C).

Our results raise a number of important questions for future studies: Does the axial elongation structure of modular networks emerge from the spatially correlated patterns of the retinal waves, or do the wave patterns build on and amplify pre-existing connectivity biases? In other words, do retinal waves play a key role in specifying the orientation identity of individual modules? Future experiments involving long-term wave blockade will be necessary to disambiguate these possibilities and to quantify the relative contributions of spontaneous wave activity versus activity-independent mechanisms to the formation of orientation-specific intrinsic connectivity.

Another open question is the role of retinal waves in establishing stable sensory responses at eye opening. In ferret, experience significantly alters visual responses after eye opening^7,25^. In contrast, our findings in tree shrews show a stable correspondence between spontaneous patterns and later evoked responses. This could reflect a species difference, where the tree shrew exhibits a more hardwired organization. However, it remains possible that experience refines certain features of the representation of orientation in the shrew post eye opening. For example, the relatively weaker retinal wave prediction of oblique vs. cardinal orientation responses may indicate that non-cardinal selectivity benefits more from sensory input following the onset of experience. But it is also worth noting that the stronger prediction of the cardinal response may reflect the more robust representation of cardinal orientations compared to oblique in developing^26^ and even adult species^27–29^.

Although retinal waves are classically associated with establishing retinotopy, our findings suggest a broader role in organizing feature-selective cortical maps. For instance, retinal waves may be driving co-activations across spatially displaced populations along an axis, reinforcing patterns of correlation that underlie modular organization. This would support the idea that orientation preference and long-range correlation structure may arise simultaneously and in an interdependent manner, rather than sequentially^30^. Our results expand our understanding of how retinal waves may be contributing to the organization of cortical structure, contributing not only to retinotopy but also to the the modular representation of orientation selectivity and to specific patterns of network connectivity that are critical for the functional alignment of both representations.

## Methods

### Animals

All experimental procedures were approved by the Max Planck Florida Institute for Neuroscience Institutional Animal Care and Use Committee and were performed in compliance with guidelines from the U.S. National Institutes of Health. Tree shrews neonates (*Tupaia belangeri*) used in this study (n = 11) were either male or female, 2-26 days old. Neonates were housed with their siblings in an incubator set at 37-38 with a 12 hour lights on/off cycle. They are hand fed each morning; procedures described below were delayed to 1 hour after feeding time.

### Viral injections

We expressed GCaMP7s by microinjecting AAV9 expressing hSyn.jGCaMP7s.WPRE (Addgene) into primary visual cortex at P2-4, ∼1 mm lateral, ∼1–2 mm anterior of lambda. Anesthesia was induced with isoflurane (2%) and maintained with isoflurane as needed based on animal vitals, but not surpassing 2%. Body temperature was maintained with a thermostatically controlled heating blanket; both rectal temperature and temperature of the heating pad surface was monitored throughout the surgery. Heart rate is monitored throughout the surgery with custom made EKG leads. Using aseptic surgical technique, skin overlying visual cortex was retracted, and a small burr hole will be made with a hand held drill (Fordom Electric Co.). Approximately 1 µL of virus contained in a pulled-glass pipette was pressure injected into the cortex at ∼200 µm below the surface over 2 min using a Nanoject-II (World Precision Instruments) in up to 3 different locations. Following the injections, skin was sutured closed and affixed with Vetbond, and the animal was recovered and returned to its incubator.

### Cranial window surgery

For experiments monitoring neuronal activity using chronic multiphoton calcium imaging approximately 10 days after initial injection procedure described above, tree shrews were retrieved from the incubator and anesthetized with 2% isoflurane. Glycopyrrolate (0.01-0.02 mg/kg) was injected subcutaneously for heart rate stabilization and minimizing secretions in the mouth and respiratory passages and lidocaine 2% mixed 1:1 with ropivacaine 0.25% was injected subcutaneously at incision site for local anesthetic effect. Animals were placed on a feedback-controlled heating pad to maintain an internal temperature of 37–38°C. Isoflurane was delivered between 1 and 2% throughout the surgical procedure to maintain a surgical plane of anesthesia., the location confirmed by A metal headplate (8mm DIA) was then implanted over V1, affixed to the skull using Metabond. The location of the headplate was confirmed by fluorescent emission of the intracranial virus injection. A craniotomy (∼5 mm diameter) was then performed and dura was removed to expose the brain, followed by an 8 mm cover glass glued into the headplate chamber. The imaging chamber was then further covered with a custom-made silicone plug (Kwik-Sil, WPI). During chronic imaging sessions, external silicone plug is carefully removed to allow for optical access and replaced by the ends of data collection sessions. Tree shrews were initially anesthetized with isoflurane (1-2%), and then reduced to low isoflurane (0.5%) to ensure for cortical activity and responses to visual stimuli. EKG and internal temperature were continuously monitored during imaging sessions. If there are any signs of distress, additional levels of anesthesia are administered immediately. Animals were then recovered and returned to their incubators.

### Calcium imaging

Wide-field epifluorescence imaging was performed using the Bergamo II Scope (Thorlabs) with blue LED illumination using an LED driver (Thorlabs). GCaMP7s fluorescence signal from the cortical surface was acquired at 10 Hz (640 × 540 pixels, field of view (FOV) with a 2x Olympus objective of 8×8 mm^2^) using a Zyla 5.5 sCMOS camera (Andor) controlled by μManager2. Individual sessions were initially registered prior to recording using a blood vessel template recorded on the first chronic session. More accurate registration was performed offline using custom scripts based on pixel correlation. Spontaneous activity was recorded in 10 minute intervals for a minimum of 1 hour per animal per imaging day, with the animal sitting in a darkened room facing an LCD monitor displaying a black screen. The animal’s heart rate was recorded for each imaging interval in order to compare anesthetic planes across experiments.

Two-photon imaging experiments were performed using the Bergamo II Scope (Thorlabs) with 920 nm excitation provided by an InSight Dual (Spectra-Physics), controlled by ScanImage software (Vidrio Technologies). Average excitation power at the exit of the objective (16 ×, CFI75, Nikon Instruments, and in 2 experiments the 10x Super Apochromatic Thorlabs objective was used) ranged from 20 to 50 mW. Images were acquired at 15 Hz (512 × 512 pixels, FOV ranges from 1.1 × 1.1 mm^2^ to 2 × 2 mm^2^. Imaging depth was between 100 and 200 um below the surface, corresponding to Layer 2/3 of the tree shrew visual cortex. Z-projections of two-photon fields of view were aligned to the epi-fluorescence imaging with the blood vessel pattern.

### Pharmacology

For acute intraocular injections during imaging sessions, isoflurane was increased to 2% for 20 minutes prior to the procedure and the eyelid was gently coaxed open. These experiments were done on p17 tree shrews. While the eye was irrigated with sterile saline, we injected 1uL of 0.9% saline or 2.5mM TTX into the vitreal humor using a glass micropipette. After injection, the needle was left in place for 30 seconds to minimize backflow upon removal. Isoflurane was decreased back to the imaging plane and cellular imagining resumed immediately. The efficacy of TTX was tested by the absence of visually evoked responses to flashing full-field luminance steps. For subcutaneous atropine injections, 1mg/kg atropine was injected subcutaneously between the shoulder blades during imaging sessions.

### Imaging processing

The region of interest (ROI) was manually drawn around the imaging window. Automated image segmentation and calcium event detection was then performed using custom routines written in MATLAB. The mean pixel intensity at each pixel location, F0 was subtracted and normalized to each frame, Ft of the movie to form a ΔF/F array: A = (Ft-F0)/F0. Each frame was smoothed with a Gaussian having a standard deviation of 250μm and a signal intensity thresholded at T, which was set to > 2 standard deviations of the signal. Calcium domain signals were automatically segmented as contiguously connected components in space and time from the resulting binarized image. A set of regional measurements were obtained from the largest component in the binary wave mask (‘area’, ‘centroid’, ‘eccentricity’, ‘majoraxislength’, ‘minoraxislength’, and ‘orientation’ from MATLAB Image Processing Toolbox). Measurements such as orientation and axis lengths were calculated by fitting an ellipse with the same second-moments as the region; the angle between the x-axis and the major axis of the ellipse was the resulting orientation which was then corrected based on the angle of the V1/V2 border when visible. Components located outside the boundaries of the region of interest or having a duration of 1 frame were ignored. The number of contiguous frames (bounding box depth) for each segmented calcium domain was taken as the wave duration.

### Pixel-based correlations of spontaneous activity patterns

For seed-based correlation mapping, pixels were selected from a 3D image stage of the spontaneous recordings where the time trace of a selected pixel was extracted and the Pearson correlation coefficient between its trace and the trace in every pixel in the ROI was computed. The correlation values were then mapped back into their original spatial locations. To compute the orientation of the resulting seed correlation map, the top 25% of non-zero values in the smoothed map results were used to define a binary mask, after which region properties of the largest detected region were extracted as described above. The orientation value of the binary mask was then corrected relative to the orientation of the V1/V2 border in the imaging field.

### Visual stimuli and Orientation fitting

Visual stimuli were delivered on a monitor using PsychoPy. The monitor had resolution 1920×1080 pixels and at the typical distance of 20 cm from the eyes extended ∼120 × 70°. The center of the monitor was matched to the mid-point between the two eyes. To evoke orientation responses, full-field square gratings at 50% contrast, 0.2 cycles per degree spatial frequency, and drifting at 4Hz temporal frequency were presented. 8 stimulus orientations were sampled. Stimuli were randomly interleaved and presented for 3s followed by 5s of gray screen. 8-10 trials of each stimulus, including blank, were shown. Orientation response maps were calculated such that each trial’s time-averaged response 1 second prior to stimulus onset (F_pre_) was subtracted from and normalized to the time averaged response across stimulus frames, F_stim_: = (F_stim_-F_pre_)/F_pre_. For fitting spontaneous correlation maps to the orientation response after eye opening, the binary mask of the spontaneous correlation was applied to the orientation response map and the orientation response value of the captured pixels was tallied. To compare results across animal maps, the resulting histograms were normalized to the maximum number of pixels in any orientation group.

### Statistics

For all comparisons in response properties across conditions and age groups we used repeated measures analysis of variance (ANOVA) followed by Tukey’s HSD post hoc test when analyzing the effects of multiple grouping factors with p<0.05 set as significance, unless otherwise noted. All graphs report means with the 95% confidence interval unless otherwise noted. Stars in figures indicate the following: ^∗^= p < 0.05, ^∗∗^= p < 0.01, ^∗∗∗^= p < 0.001. Values of n are indicated in the figure legends. Custom code was written using Matlab R2019b. All code is available upon request.

## Acknowledgements

We would like to thank J. Kerr, G. Kreal, R. Satterfield and N. Shultz for technical assistance, as well as members of the Fitzpatrick laboratory for helpful discussions. This research was supported by US National Institutes of Health grant EY006821 as well as the Max Planck Florida Institute for Neuroscience.

## Author contributions

D. F. and A. G. contributed to experimental design, result interpretation, and writing of the manuscript. A.G. performed all experiments and data analysis.

## Author information

The authors declare no competing financial interests. Correspondence and requests for materials should be addressed to D.F.

